## Supplementary material for "Decoding brain states on the intrinsic manifold of human brain dynamics across wakefulness and sleep"

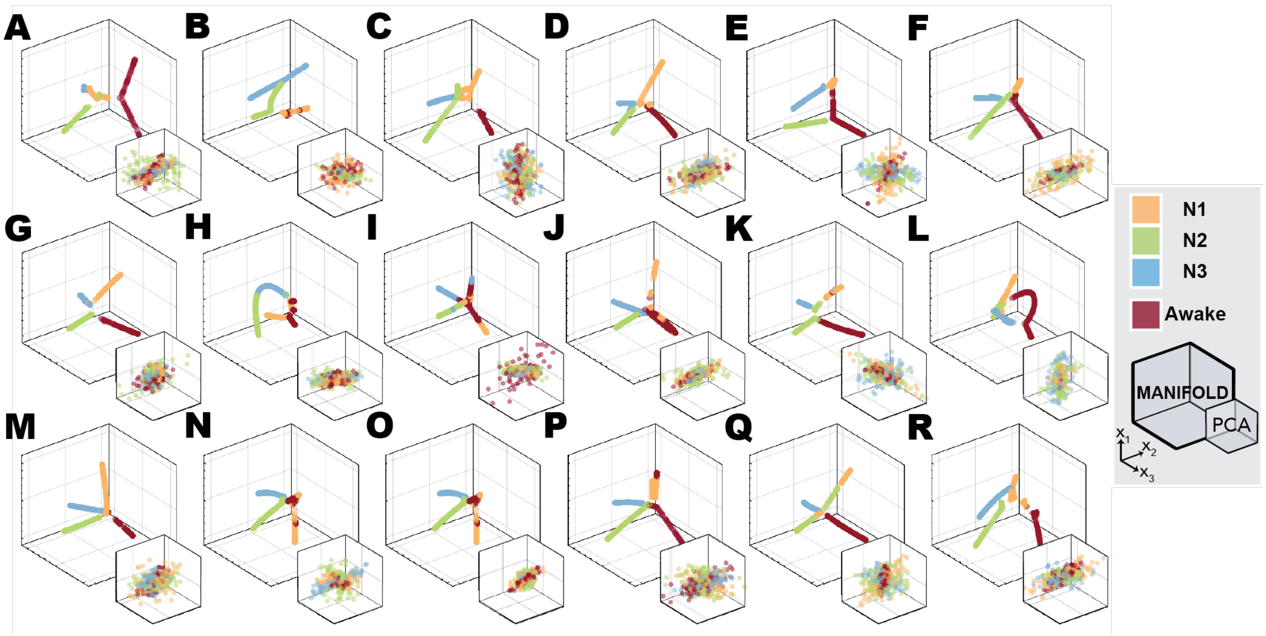

***Fig. S1**. **Representation of the individual subjects' fMRI BOLD data during wakefulness and sleep embedded into the lower dimensional spaces***. The plots show the fMRI BOLD data embedded into the three first dimensions of the *intrinsic manifold* (large coordinate system) and into the three principal components derived from PCA (small coordinate system shown at each corner). Each separate coordinate system corresponds to the data of eighteen different participants, embedded individually.

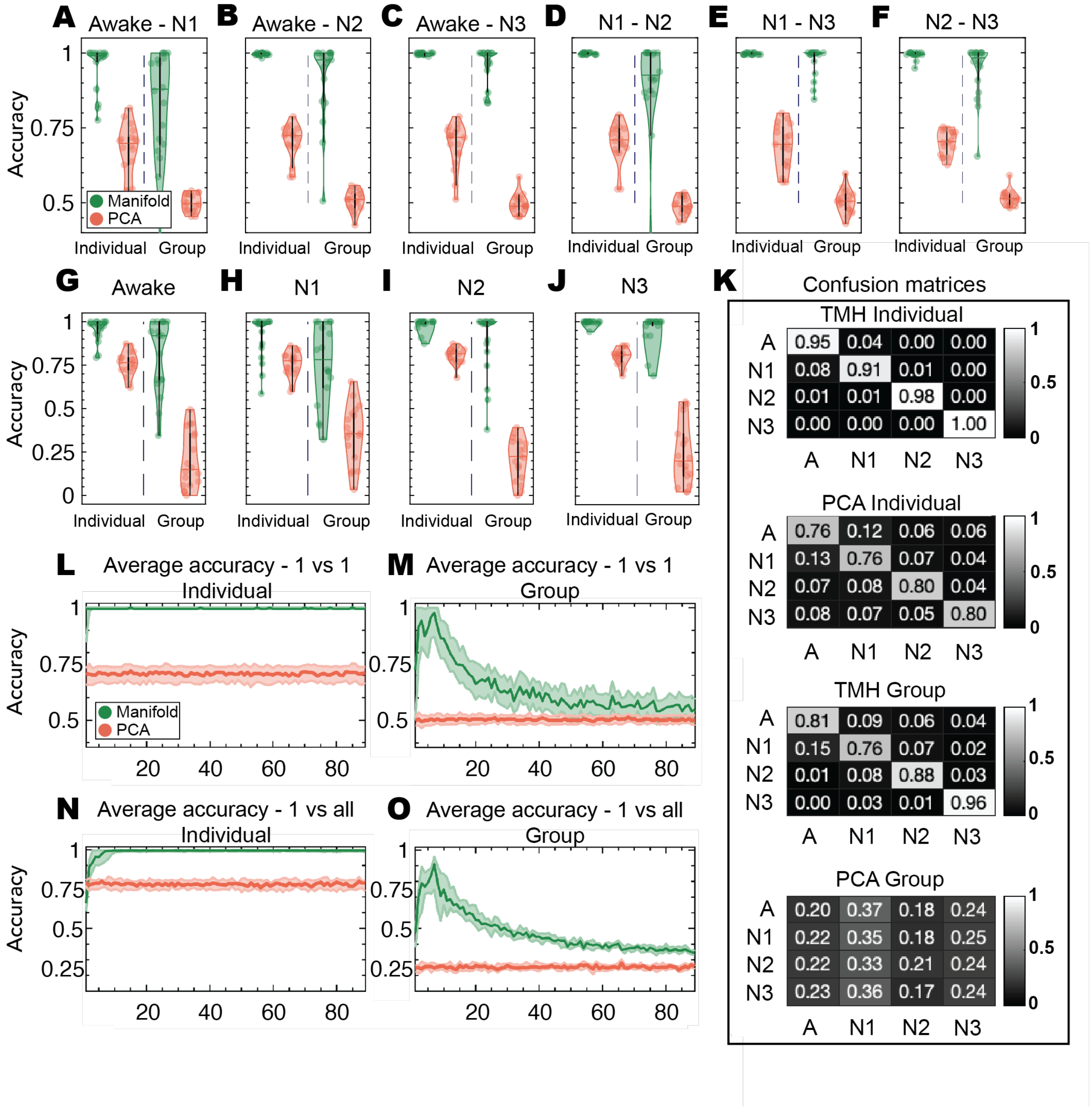

**Fig. S2. Accuracy of brain state decoding on the intrinsic manifold of brain dynamics and on PCA for 3 dimensions using a linear SVM. A-F)** The accuracies of the SVM 1-vs-1 classification between **A)** wakefulness and N1, **B)** wakefulness and N2, **C)** wakefulness and N3, **D)** N1 and N2, **E)** N1 and N3, and **F)** N2 and N3. **G-J)** The accuracies of the SVM 1-vs-all classification for each stage: **G)** wakefulness, **H)** N1, **I)** N2 and **J)** N3. The accuracy is defined as the ratio between the number of true positives and the total number of tested time points. The boxplots' centrality is indicated by the median, and the boxes extend between 25-*th* and 75-*th* percentiles. Each colored circle corresponds to the classification accuracy for each single subject (in the case of individual analysis, left of the central dashed line) and to the accuracy of each leave-one-subject-out round (in the case of group analysis, right to the central dashed line). The classification accuracies on the intrinsic manifold and in PCA space are shown in green and red dots, respectively. Classifications are performed in spaces of dimensionality d=3. For all classifications, intrinsic manifold classification yields significantly better accuracies (for all comparisons, *p-value*<.001, Wilcoxon Rank-sum two-sided test, corrected for multiple comparisons via FDR). **K)** Confusion matrices obtained from the 1-vs-all classification experiments (shown in **G-J)**). **L-M)** show the average accuracy across all stage-to-stage classifications for varying dimensionality of the embedding spaces for individual participants (**L)**) and for group analysis (**M)**), respectively. **N-O)** show the average accuracy across all stage (1-vs-all) classifications for varying dimensionality of the embedding spaces for individual participants (**N)**) and for group analysis (**O)**), respectively. The solid lines indicate the median of the distribution across classifications and shaded areas indicate 25-*th* and 75-*th* percentiles.

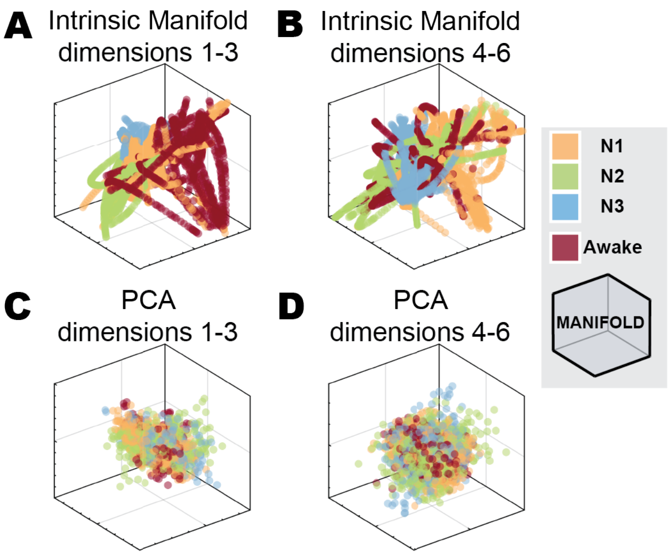

**Fig. S3. Representation of the fMRI BOLD data during wakefulness and sleep at the group level embedded in lower dimensional spaces**. The plots show the intrinsic manifolds aligned to a common reference for 18 participants. The six first dimensions of the intrinsic manifold (big coordinate system) and into the six principal components derived from PCA (small coordinate system). For all cases, nonlinear embedding of the data into their intrinsic manifold led to well-structured intrinsic manifolds with a clearer separation of different sleep stages (as defined through polysomnography) compared to the PCA linear embedding.

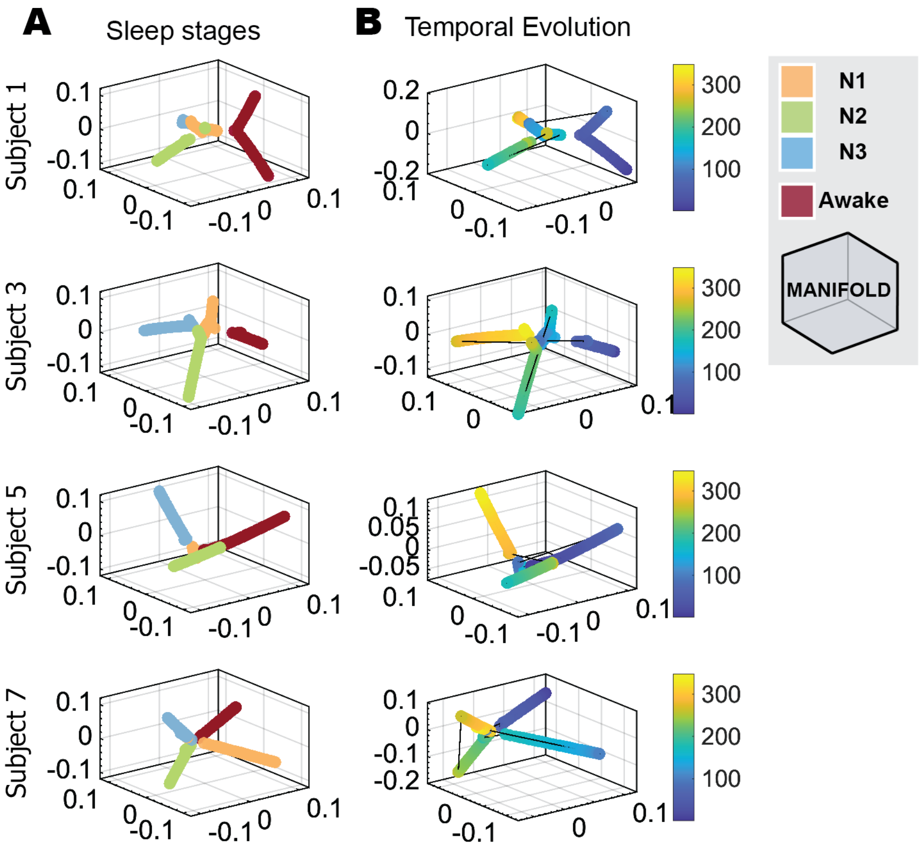

**Fig. S4. Temporal evolution of the fMRI BOLD data during wakefulness and sleep embedded in lower dimensional spaces**. The plots show the intrinsic three-dimensional manifolds of four subjects, with color coding for both (A) sleep stage and (B) time-index. For all subjects, fMRI BOLD data shows smooth intra-stage transitions and inter-stage shortcuts.

**Table S1. SVM accuracy medians comparison through *Wilcoxon Ranksum* two-sided test. *p-values* corrected for multiple comparisons via FDR.** All reported accuracies for the SVM classification are higher in the intrinsic manifold, and significant differences are marked with asterisks (** $p$<.005, * $p$<.05, Monte-Carlo phase-randomized simulations, corrected for multiple comparisons via FDR).

| | *Individual manifold* | | | *Group manifold* | | |
|---|---|---|---|---|---|---|
| | Intrinsic manifold accuracy | PCA accuracy | Median comparison $p$-value | Intrinsic manifold accuracy | PCA accuracy | Median comparison $p$-value |
| *1 vs 1* | 0.99±0.03 ** | 0.69±0.07 ** | <.001 | 0.92±0.13 ** | 0.50±0.03 | <.001 |
| *1 vs all* | 0.96±0.04 ** | 0.78±0.03 ** | <.001 | 0.85±0.09 ** | 0.25±0.07 | <.001 |

**Table S2. SVM accuracy medians comparison for each class, through *Wilcoxon Ranksum* two-sided test. *p-values* corrected for multiple comparisons via FDR.** All reported accuracies for the SVM classification are higher in the intrinsic manifold, and all differences are significant (** $p$<.005, * $p$<.05, Ranksum two-sided test, corrected for multiple comparisons via FDR).

| | | *Individual manifold* | | | *Group manifold* | | |
|---|---|---|---|---|---|---|---|
| | | Intrinsic manifold accuracy | PCA accuracy | Median comparison *p*-value | Intrinsic manifold accuracy | PCA accuracy | Median comparison *p*-value |
| *1 vs 1* | *Awake – N1* | 0.96±0.07 | 0.67±0.09 | ** | 0.82±0.20 | 0.50±0.03 | ** |
| | *Awake – N2* | 0.99±0.01 | 0.71±0.06 | ** | 0.91±0.14 | 0.51±0.03 | ** |
| | *Awake – N3* | 0.99±0.01 | 0.69±0.07 | ** | 0.96±0.06 | 0.49±0.03 | ** |
| | *N1 – N2* | 0.99±0.01 | 0.70±0.07 | ** | 0.90±0.15 | 0.49±0.03 | ** |
| | *N1 – N3* | 0.99±0.01 | 0.69±0.07 | ** | 0.97±0.05 | 0.51±0.04 | ** |
| | *N2 – N3* | 0.99±0.01 | 0.70±0.04 | ** | 0.94±0.09 | 0.52±0.03 | ** |
| *1 vs all* | *Awake* | 0.95±0.07 | 0.76±0.07 | ** | 0.81±0.21 | 0.20±0.16 | ** |
| | *N1* | 0.91±0.13 | 0.76±0.07 | ** | 0.76±0.23 | 0.35±0.16 | ** |
| | *N2* | 0.98±0.03 | 0.80±0.05 | ** | 0.88±0.19 | 0.21±0.12 | ** |
| | *N3* | 0.99±0.01 | 0.80±0.05 | ** | 0.96±0.09 | 0.24±0.18 | ** |

**Table S3. AUC medians comparison through *Wilcoxon Ranksum* two-sided test. p-values corrected for multiple comparisons via FDR.** All AUC are significantly higher for the intrinsic manifold than for the projection into the PCA space.

| | *Individual manifold* | | |
|---|---|---|---|
| | Intrinsic manifold AUC | PCA AUC | Median comparison *p*-value |
| *Awake – N1* | 0.99±0.04 | 0.51±0.01 | <.001 |
| *Awake – N2* | 0.98±0.04 | 0.51±0.01 | <.001 |
| *Awake – N3* | 0.99±0.04 | 0.51±0.01 | <.001 |
| *N1 – N2* | 0.99±0.04 | 0.51±0.01 | <.001 |
| *N1 – N3* | 1.00±0.01 | 0.51±0.01 | <.001 |
| *N2 – N3* | 0.94±0.10 | 0.52±0.02 | <.001 |

**Table S4. Decoding accuracy compared to *Tagliazucchi 2012 Neuroimage*.** This table reports the results of *Tagliazzuchi et al. Neuroimage 2012,* in comparison to our results on group manifolds, and on individual manifolds. We believe that the comparison between our accuracies on individual manifolds and previous efforts is the fairest comparison, as in both these cases, data from the same subjects are included in both training and testing sets.

| | Tagliazucci 2012 (TW: 1 min; **max accuracy** out of 6-fold – see their TABLE 5) | Intrinsic manifold (GROUP, single time-points; **mean accuracy** out of leave-one-subject-out) | Intrinsic manifold (INDIVIDUAL, single time-points; **mean accuracy** out of 6-fold cross-validation) |
|---|---|---|---|
| *Awake – N1* | 0.76 | 0.82 | **0.96** |
| *Awake – N2* | 0.88 | 0.91 | **0.99** |
| *Awake – N3* | 0.89 | 0.96 | **0.99** |
| *N1 – N2* | 0.83 | 0.90 | **0.99** |
| *N1 – N3* | 0.93 | 0.97 | **0.99** |
| *N2 – N3* | 0.87 | 0.94 | **0.99** |

**Video S5. Temporal evolution of the fMRI BOLD data during wakefulness and sleep embedded in lower dimensional spaces for one subject.** The video shows the intrinsic three-dimensional manifold of subject 7, with color coding for both (A) sleep stage and (B) time-index. fMRI BOLD data shows smooth intra-stage transitions and inter-stage shortcuts. The frame rate is 10 times faster**.**
